# The Unique Neural Signature of Your Trip: Functional Connectome Fingerprints of Subjective Psilocybin Experience

**DOI:** 10.1101/2023.03.20.532894

**Authors:** Hanna M. Tolle, Juan Carlos Farah, Pablo Mallaroni, Natasha L. Mason, Johannes G. Ramaekers, Enrico Amico

## Abstract

The emerging neuroscientific frontier of brain fingerprinting has recently established that human functional connectomes (FCs) exhibit *fingerprint-like* idiosyncratic features, which map onto heterogeneously distributed behavioural traits. Here, we harness brain-fingerprinting tools to extract FC features that predict subjective drug experience induced by the psychedelic psilocybin. Specifically, in neuroimaging data of healthy volunteers under the acute influence of psilocybin or a placebo, we show that, post psilocybin administration, FCs become more idiosyncratic due to greater inter-subject dissimilarity. Moreover, whereas in placebo subjects idiosyncratic features are primarily found in the frontoparietal network, in psilocybin subjects they concentrate in the default-mode network (DMN). Crucially, isolating the latter revealed an FC pattern that predicts subjective psilocybin experience and is characterised by reduced within-DMN and DMN-limbic connectivity, as well as increased connectivity between the DMN and attentional systems. Overall, these results contribute to bridging the gap between psilocybin-mediated effects on brain and behaviour, while demonstrating the value of a brain-fingerprinting approach to pharmacological neuroimaging.

**Author summary:** The trending field of brain fingerprinting focuses on characterising fingerprint-like idiosyncratic features of human functional connectomes (FCs), which have been shown to predict heterogeneously distributed behavioural traits. Here, we apply brain-fingerprinting methods to fMRI data from subjects who were administered the psychedelic psilocybin or a placebo. We find that, compared to the placebo condition, subjects under acute psilocybin effects exhibited more idiosyncratic FCs, with idiosyncratic features being largely concentrated in the default-mode network (DMN). Furthermore, we isolated an idiosyncratic FC pattern that predicted reports of subjective psilocybin experiences. This pattern was characterised by altered DMN connectivity, specifically by reduced within-DMN and DMN-limbic connectivity, and increased connectivity between the DMN and attentional systems. This work paves the way for exciting new research harnessing pharmacological brain fingerprinting.

## Introduction

Psychedelic drugs such as lysergic acid diethylamide (LSD), dimethyltryptamine (DMT), and psilocybin are well-known for their ability to transiently yet profoundly alter an individual’s perception of oneself and the external world, inducing what is called an altered state of consciousness (ASC) (Studerus et al., 2010). Despite their long history of medicinal and ceremonial use, especially in indigenous contexts, psychedelics have only recently been rediscovered (R. L. Carhart-Harris & Goodwin, 2017) as putatively effective and safe (Johnson et al., 2018, 2019) treatments for a range of psychiatric disorders. Most notably, psilocybin-assisted psychotherapy has been reported to yield significant improvements of clinical symptoms in major depressive disorder (Daws et al., 2022; Rucker et al., 2016), obsessive compulsive disorder (Moreno et al., 2006), addiction (Bogenschutz et al., 2015; Johnson et al., 2017), and end-of-life distress (Griffiths et al., 2016; Ross et al., 2016).

There is strong evidence that the characteristic mind-altering effects of psychedelics are primarily mediated via agonism of the serotonin receptor 5-HT2AR (Glennon et al., 1992; Stenbæk et al., 2021; Vollenweider et al., 1998). Nevertheless, how 5-HT2AR agonism affects whole-brain dynamics to induce the experienced ASC is still a subject of ongoing research. Major contributions to this end have been made by network neuroscience-based approaches, examining psychedelic-induced changes of the brain’s functional connectome (FC) architecture (R. L. Carhart-Harris et al., 2013a). The FC is a statistical model representing the brain as a network composed of brain regions (nodes) and the connections thereof (edges), where the connection between two brain regions typically encodes the Pearson correlation of their neurophysiological time series (Bullmore & Sporns, 2009). In functional magnetic resonance imaging (fMRI), a salient hallmark of the human FC is the strongly correlated hemodynamic activity within distinct groups of brain regions, giving rise to the so-called resting-state networks (RSNs) (Yeo et al., 2011). Regions of the same RSN are believed to collaborate in serving specific functions that require increasingly complex information processing, ranging from unimodal sensory regions of the somatomotor (SM) and visual (Vis) networks to heteromodal association cortices of the frontoparietal (FPN) and default-mode (DMN) networks (Luppi et al., 2022; Margulies et al., 2016).

In this context, it has been proposed that the primary action of psychedelics, whereby the subjective effects are elicited, is to reconfigure the brain’s functional hierarchy by modulating the activity of higher-level cortical networks (R. L. Carhart-Harris & Friston, 2019; Corlett et al., 2019; Doss et al., 2021), which are known to exhibit exceptionally high 5-HT2AR densities (Beliveau et al., 2017). Indeed, reduced activity and functional connectivity within these higher-level systems, in particular the DMN, are among the most frequently reported neural correlates of the acute psychedelic state, and have been shown to correlate with the intensity of subjective effects (R. L. Carhart-Harris et al., 2012, 2016; Madsen et al., 2021; Muthukumaraswamy et al., 2013; Smigielski et al., 2019). Concomitantly, another consistently observed acute neural response to psychedelic drugs is an increase in functional connectivity between lower-level systems (Preller et al., 2018, 2020).

While network-based analyses of neuroimaging data have advanced our understanding of the group-level effect of psychedelics on the functional network organisation of the brain, it is less clear how each individual’s neural response to a psychedelic compound maps onto the various aspects of their unique subjective drug experience. In fact, the subjective effects of psychedelics are multifaceted and highly variable, both within and between subjects (Moujaes et al., 2022). Crucially, several lines of evidence suggest that the long-term benefits of psilocybin are influenced by the subjective acute drug experience (Bogenschutz et al., 2015; Griffiths et al., 2016; Johnson et al., 2017; Roseman et al., 2018; Ross et al., 2016). This emphasises the need for further research investigating how an individual’s unique psychedelic experience emerges from the underlying FC reconfigurations in that person’s brain.

Mapping FC changes onto subjective drug-experience reports is a non-trivial task and one major reason for this is the high dimensionality of the FC, comprising the edge weights of all unique pairs of brain regions, in contrast to the typically low sample sizes of neuroimaging studies. Intriguingly, seminal work from Finn *et al*. (Finn et al., 2015) revealed that the weights of specific FC edges are idiosyncratic, i.e. they are both unique to each subject and reliable across test-retest fMRI scanning sessions, just like the pattern of furrows and ridges that defines an individual’s fingerprint. Importantly, particularly idiosyncratic edges that facilitate the identification of subjects were also shown to be most predictive of behavioural traits (Finn et al., 2015). These findings have inspired a series of other studies focusing on idiosyncratic rather than generalisable FC patterns, giving rise to the trending field of brain fingerprinting (Amico & Goñi, 2018; Mallaroni et al., 2022; Stampacchia et al., 2022; Van De Ville et al., 2021).

In this work, we propose that neuroimaging studies investigating the effects of pharmacological substances on brain activity stand to gain significantly from a *pharmacological brain-fingerprinting* approach that integrates brain-fingerprinting methods. We expect this approach to be especially illuminating if individual responses to the drug in question are heterogeneous. Notably, the acute subjective experience of psychedelic compounds is highly variable across individuals (Moujaes et al., 2022) and has been associated with long-term drug effects (Nichols, 2020). This renders pharmacological brain fingerprinting a particularly promising approach for studying individual drug responses to psychedelics using neuroimaging data, with great potential to inform precision medicine.

In this vein, a pioneering pharmacological fingerprinting study reported that, under the influence of the psychedelic compound ayahuasca, FC fingerprints are altered such that they predict aspects of individual psychedelic experience (Mallaroni et al., 2022). However, to the best of our knowledge, no study to date has investigated the acute effect of psilocybin on FC fingerprints and its relevance to subjective psilocybin experience.

Addressing these knowledge gaps, this study applies brain-fingerprinting tools to 7-Tesla fMRI data of 46 healthy volunteers who were administered 0.17 mg/kg psilocybin (n=21) or a placebo (n=25), in order to i) compare FC fingerprints between psilocybin and placebo conditions, and ii) examine whether idiosyncratic FC patterns allow for more accurate predictions of subjective psilocybin experience with respect to FC patterns composed of randomly selected features. Our measure of subjective psilocybin experience captured 11 aspects of the unique psilocybin-induced ASC of each subject, which were assessed retrospectively using the well-established 5D-ASC questionnaire (Dittrich, 1998; Studerus et al., 2010).

Interestingly, we find that in the psilocybin group FCs are significantly more dissimilar and thus allow for better subject identification than in the placebo group. Furthermore, we present a simple heuristic for estimating the idiosyncrasy of single FC edges, revealing that in placebo idiosyncratic edges are primarily located in FPN regions such as the lateral prefrontal cortex (PFC), whereas in psilocybin they concentrate in DMN regions such as medial prefrontal (mPFC), posterior cingulate (PCC) and inferior parietal (IPC) cortices. Finally, by selecting highly idiosyncratic edges we uncovered an FC pattern that significantly predicts subjective psilocybin experience, characterised by decreased within-DMN and DMN-limbic connectivity, and increased connectivity between DMN and attentional networks.

## Results

### Introducing drug brain fingerprinting

Step one of our *pharmacological brain-fingerprinting* approach is the derivation of two FCs per subject from the first and second temporal halves of the regional fMRI blood-oxygen-level-dependent (BOLD) time series (Figure 1A). These two FCs, which we will respectively refer to as first-half and second-half FC, are used to assess the reliability of FC features. Reliability in our analysis should therefore be interpreted in terms of the temporal stability of an individual’s FC pattern. Although related, this differs from the notion of test-retest reliability where repeated measurements are taken during separate acquisitions, so as to account for session-to-session variability due to, for instance, methodological differences.

**Figure 1.**
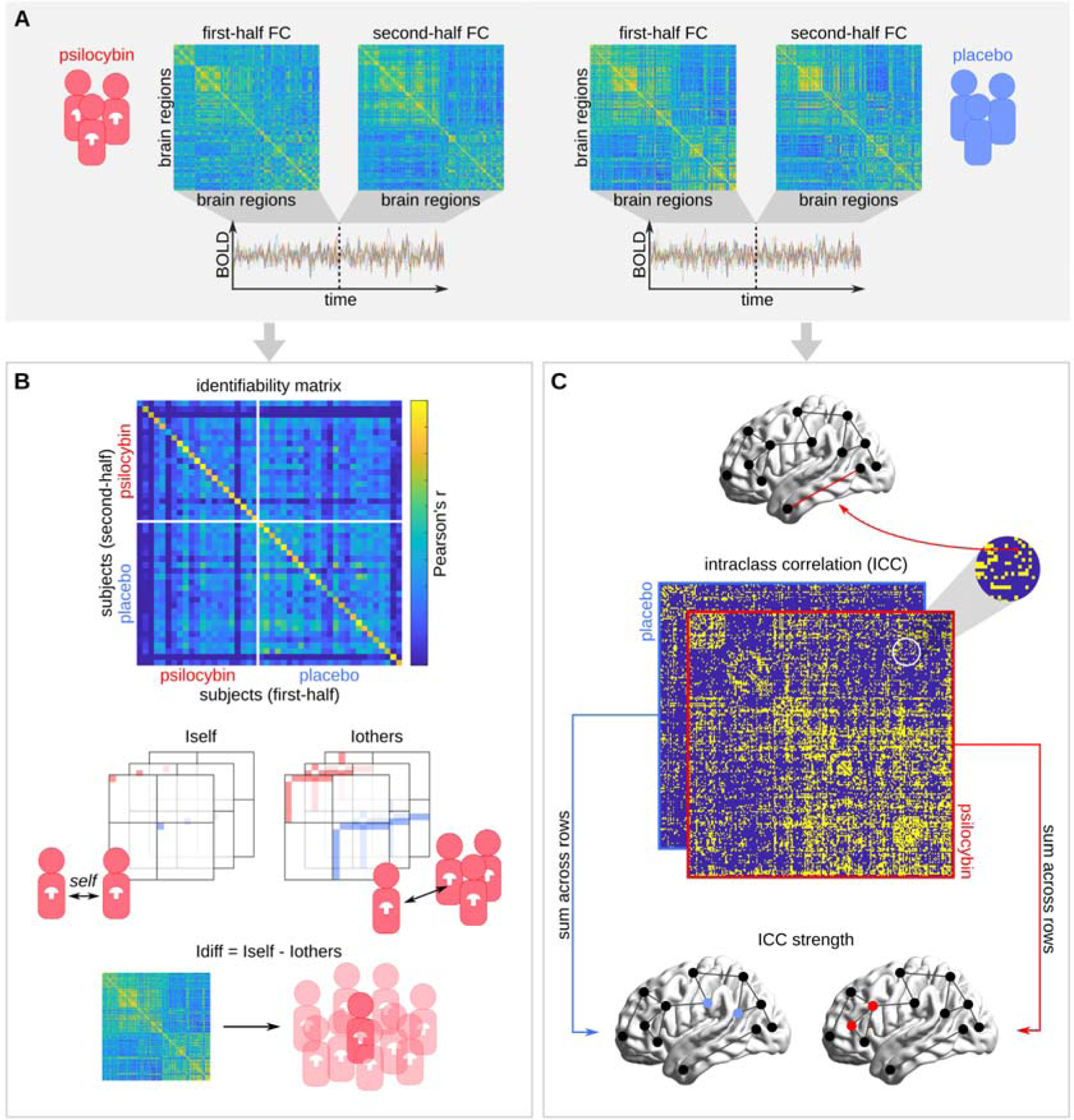
Pharmacological brain-fingerprinting workflow. (A) The first-half and second-half FCs of each subject are computed, respectively, from the first and second half of the subject’s regional fMRI BOLD time series such that the resulting symmetric square matrices encode the Pearson correlation of the BOLD signals of each unique pair of brain regions during the first half of the scan (first-half FC) and the second half of the scan (second-half FC). (B) The identifiability matrix displays the Pearson correlation between the second-half (rows) and first-half (columns) FCs of all subjects, where rows and columns are sorted such that subjects of the same condition are clustered together. Each diagonal element of the identifiability matrix encodes the *Iself* of one subject. The *Iothers* of each subject are computed by averaging over all off-diagonal elements that correspond to first-half versus second-half FC correlations between that subject and every other subject of the same condition. Finally, *Idiff*, which is defined as the difference between *Iself* and *Iothers*, indicates the within-group identifiability of a subject based on their FC. (C) For each condition, we computed the ICC of each FC edge as a measure of *idiosyncrasy*. ICC matrices are displayed as binarised, thresholded (ICC>0.6) matrices for better visualisation. The idiosyncrasy of each brain region was assessed using ICC strength, defined as the sum of non-thresholded ICC values of all edges of that brain region.

Furthermore, we adopt two different toolboxes from brain fingerprinting for i) measuring the subject identifiability based on a given set of FC edges (Figure 1B), and ii) mapping edgewise and brain-regional distributions of idiosyncrasy (Figure 1C).

Subject identifiability estimates how well a subject can be identified within a group of other subjects based on a selection of FC edges. It can be measured using *Idiff*, which is the difference between *Iself* and *Iothers* (Amico & Goñi, 2018), where *Iself* and *Iothers* respectively capture the reliability and uniqueness of the edge selection in terms of within- and between-subject FC similarity (Pearson correlation) (Figure 1B).

Idiosyncrasy intuitively describes the property of being characteristic of an individual. Consistent with this intuition, we refer to edges that promote the identification of subjects as idiosyncratic. For each condition, we measure the edgewise idiosyncrasy using intraclass correlation (ICC), and brain-regional idiosyncrasy using ICC strength, which we define to be the total ICC of all edges corresponding to a given brain region (Figure 1C).

### FC-based subject identifiability

The identifiability metrics of the full FCs of 21 psilocybin and 25 placebo subjects were, respectively, derived from the upper-left and lower-right quadrants of the identifiability matrix shown in Figure 2A. Interestingly, while there was no significant group difference in *Iself* (df=43, t-stat=-0.20, p=0.8460), psilocybin subjects tended to have lower *Iothers* (df=43, t-stat=-3.33, p=0.0018) and higher *Idiff* (df=43, t-stat=2.10, p=0.0414) than placebo subjects, suggesting that there were stable and substantial inter-individual differences in the neural response to psilocybin such that subject identification was facilitated. All reported differences were corrected for head motion as a potential confounder, which was slightly but significantly higher in the psilocybin group (df=44, t-stat=2.06, p=0.0456; see methods).

**Figure 2.**
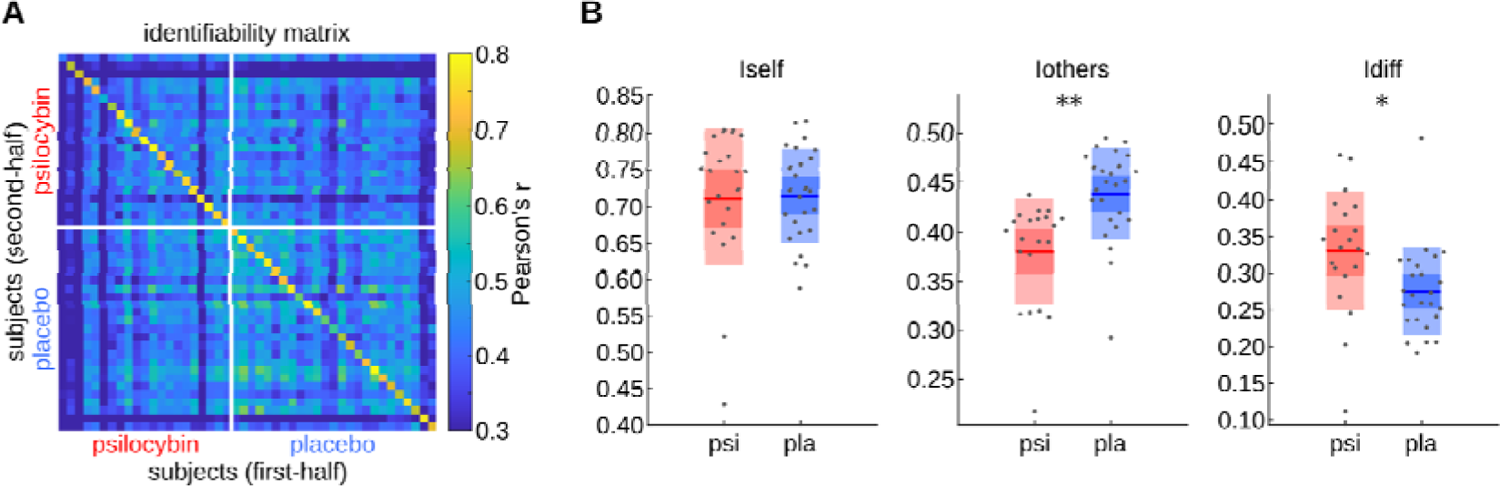
Improved identifiability in the psilocybin group due to greater FC dissimilarity. (A) Identifiability matrix as described in Figure 1B. (B) Distributions of *Iself*, *Iothers*, and *Idiff* in psilocybin (psi) and placebo (pla) conditions. The boxes display the mean, standard deviation, and 95% confidence interval of each distribution with the jittered raw data plotted on top. Significant group effects according to an ANOVA for predicting the measure from group and motion (see methods) are labelled with asterisks (* p<0.05; ** p<0.01).

### Edgewise and brain-regional distributions of idiosyncrasy

For an initial visual assessment of the spatial distribution of idiosyncrasy, we plotted the ICC of each edge (thresholded at 0.6 for better visualisation) within the psilocybin and placebo group as symmetric NxN matrices, where N=200 is the number of brain regions (Figure 3A). We will hereafter denote the non-thresholded psilocybin and placebo ICC matrices as ICCpsi and ICCpla, respectively. Figure 3B displays the difference between ICCpsi and ICCpla averaged across RSN connections. As can be seen, Vis and DMN edges tended to be more idiosyncratic in psilocybin compared to placebo, whereas the opposite trend was observed for FPN edges.

**Figure 3.**
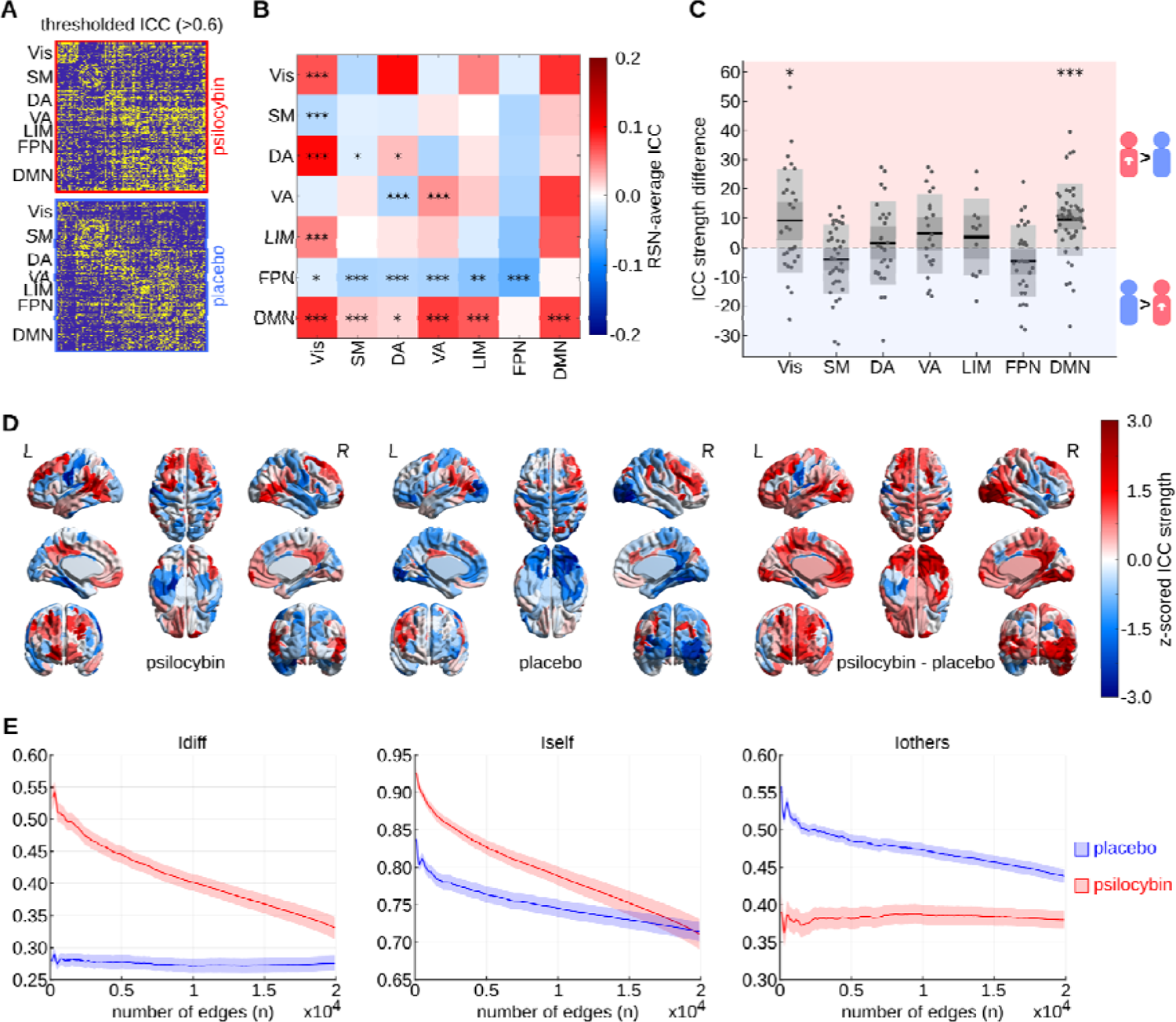
Shifted spatial distributions of idiosyncrasy in psilocybin versus placebo. (A) Thresholded (>0.6) edgewise ICC matrices of psilocybin (red) and placebo (blue) groups. (B) For each unique pair of cortical RSNs, we compared the ICCs of all edges connecting the two RSNs between psilocybin and placebo conditions. The heatmap displays the resulting mean ICC differences such that positive (red) values correspond to a higher average ICC in the psilocybin group, and vice versa. FDR-corrected significant p-values according to a paired *t*-test are highlighted in the lower triangular of the symmetric heatmap (* p<0.05; ** p<0.01; *** p<0.001). (C) Group differences in ICC strength within each RSN. Each box displays the mean, standard deviation, and 95% confidence interval of the psilocybin-placebo differences in ICC strength of all brain regions within one RSN. Distributions with means that are significantly different from 0 according to a paired *t*-test and post FDR-correction are labelled with asterisks (* p<0.05; ** p<0.01; *** p<0.001). (D) Brain renderings of z-scored ICC strength in psilocybin and placebo groups, and z-scored psilocybin minus placebo ICC-strength differences. (E) Group-average and corresponding standard error of *Idiff*, *Iself*, and *Iothers* in psilocybin (red) and placebo (blue) computed for the *n* = {50, 100, …, 19900} top ICCpsi-ranked edges. Note that selecting edges based on high ICCpsi results in an optimization of *Idiff* in the psilocybin, but not the placebo group, suggesting a functional reconfiguration of the FC fingerprint upon psilocybin administration. Vis = visual network; SM = somatomotor network; DA = dorsal attention network; VA = ventral attention network; LIM = limbic system; FPN = frontoparietal network; DMN = default-mode network.

Concurrently, we found that in the psilocybin group ICC strength was significantly higher in Vis (df=45, t-stat=2.77, p=0.0099, fdr=0.0351) and DMN (df=45, t-stat=5.25, p=0.0000, fdr=0.0000) regions and lower in FPN (df=45, t-stat=-2.07, p=0.0472, fdr=0.1000) regions compared to placebo, although the FPN effect did not survive FDR correction for seven comparisons (Figure 3C). Moreover, brain renderings of the z-scored ICC strength in placebo, psilocybin, and their difference reveal a systematic reconfiguration of the brain fingerprint: whereas in placebo idiosyncratic edges accumulate in FPN-associated right lateral prefrontal regions, in psilocybin they are primarily localised in DMN-associated regions including the mPFC, PCC, and IPC (Figure 3D).

Concurrently, we demonstrate that selecting idiosyncratic edges based on ICCpsi results in an optimization of the *Idiff* of psilocybin but not placebo subjects (Figure 3E), and the vice versa for edge selection based on high ICCpla (supplementary Figure 1). More specifically, in an iterative process we computed *Idiff*, *Iself* and *Iothers* for all subjects considering only the top *n* edges with the highest ICCpsi with *n* = {50, 100, …, 19900}. That is, the final step included all edges. As expected, the group-average *Idiff* of psilocybin subjects peaks at a low number of included edges (*n*=100) before decreasing gradually with increasing *n*, as *Iself* declines and *Iothers* slightly inclines. Remarkably however, the group-average *Idiff* of placebo subjects remains virtually flat across all values of *n*, which is despite higher *Iself* and due to higher *Iothers* in edges with high ICCpsi. This implies that temporally stable edges, which tend to be similar across individuals under normal (placebo) conditions, are differentially altered by psilocybin such that they follow a more heterogeneous distribution in psilocybin subjects.

### Idiosyncrasy-informed edge selection improves predictions of subjective psilocybin experience

Our previous results prompted the question whether idiosyncrasy (ICCpsi) can be used as a selection criterion for edges that predict subjective psilocybin experience. To this end, we first ran principal component analysis (PCA) on the behavioural data describing the individual drug experience of each psilocybin subject in 11 dimensions, which was acquired using the 5D-ASC questionnaire (Dittrich, 1998; Studerus et al., 2010) (Figure 4A). This enabled us to extract the maximally heterogeneous components in the behavioural data and reduce the dimensionality of our prediction target. The resulting first principal component (PC) explained 45.7% of the variance and the four most important behavioural aspects were “insightfulness”, “experience of unity”, “blissful state”, and “changed meaning of percepts” (Figure 4B). Notably, all PC coefficients were positive. Thus, the subject scores of this PC, henceforth denoted as *b*, may be interpreted as capturing the overall intensity of the drug experience.

**Figure 4.**
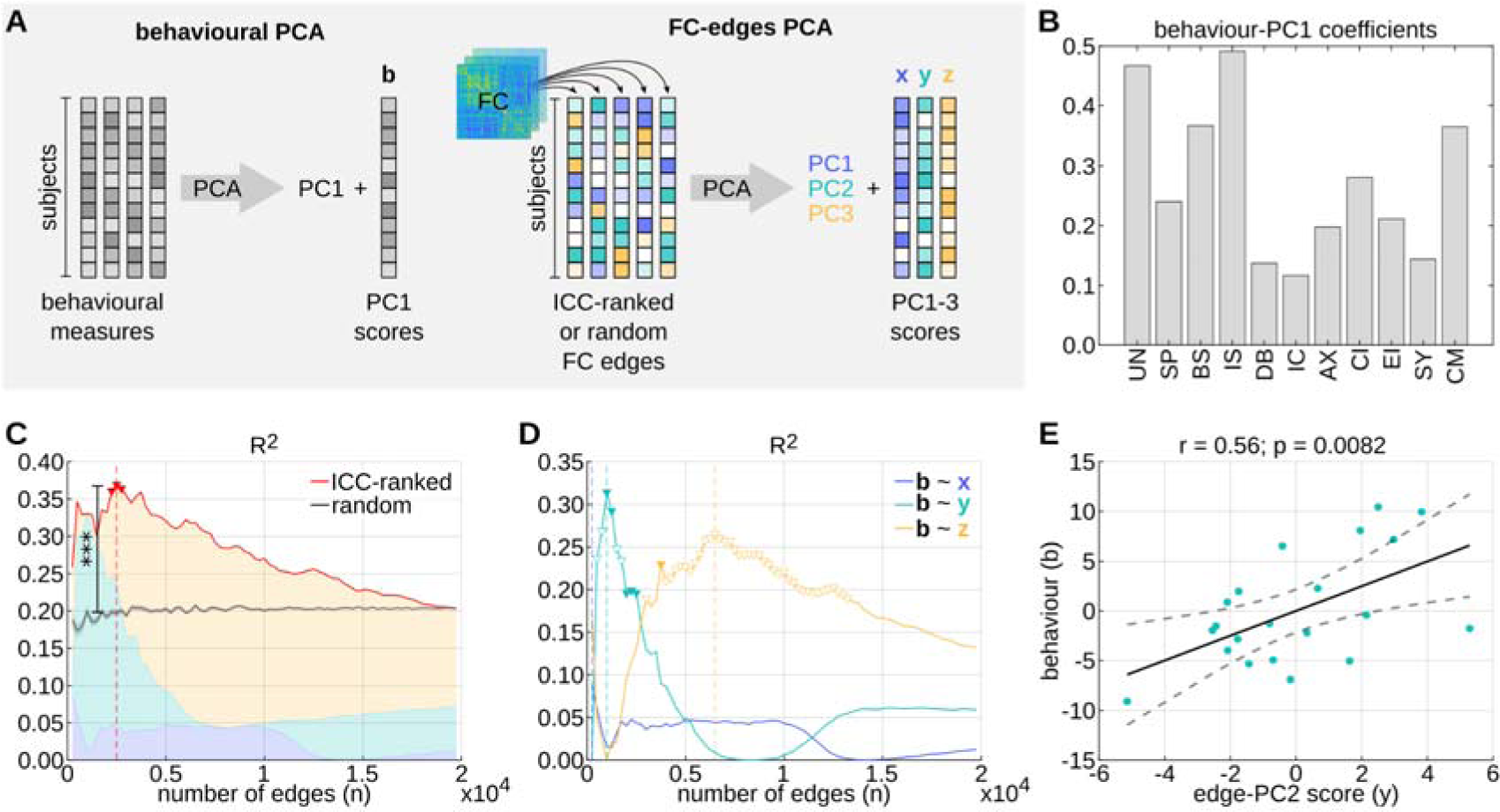
Idiosyncrasy-informed feature selection improves predictions of subjective drug experience. (A) Schematic workflow for constructing linear models to predict subjective psilocybin experience from FC edges. First, PCA was applied to the behavioural data, containing measurements of 11 aspects of subjective psilocybin experience from each subject.

Following the behavioural PCA, we applied additional PCAs to different selections of *n* edges from the FCs that were derived from the full-length BOLD time series of each psilocybin subject. Specifically, PCAs were run on i) the *n* top ICCpsi-ranked edges, and ii) 100 random selections of *n* edges from all psilocybin subjects, where *n* = {250, 500, …, 19900} (Figure 4A). Finally, for each edge selection and each *n* we built a linear model to predict *b* from the subject scores *x*, *y*, *z*, corresponding to the multidimensional FC patterns that are represented by the first three PCs (PC1, 2, 3) of the FC-edge PCA, respectively. More details on this method and the rationale behind it are provided in the methods. Remarkably, the R² value of idiosyncrasy-informed models was consistently higher than the average R² value of 100 models based on random edge selections for all *n* smaller than a threshold at which the two model types inevitably converged (Figure 4C). Moreover, a two-sided, one-sample *t*-test comparing the R² values of the models based on idiosyncratic versus randomly-selected edges at *n*=2500, where the R² value of idiosyncrasy-informed models was found to be maximal, yielded a highly significant result (p=1.00e-6, FDR-corrected for 79 steps in *n*).

In contrast to randomly selected edges, idiosyncratic edges tend to connect brain regions that are spatially close to each other and, concomitantly, more likely to have correlated BOLD time series (see Figure 3D). Thus, to ensure that the predictive advantage of idiosyncrasy-informed models was not due to the spatial contiguity of edges, we repeated the analysis from Figure 4C with spatial-contiguity-preserving surrogate models. More specifically, at each step *n* we built 100 linear models that were based on *n* edges connecting the same brain regions as the *n* most idiosyncratic edges, but the assignments of BOLD time series to each brain region were shuffled by randomly rotating spherical projections of these assignments, using software provided by Váša and Mišić (Váša & Mišić, 2022). Notably, a two-sided, one-sample *t*-test comparing the R² values of the two model types at *n*=2500 revealed that idiosyncrasy-informed models continue to outperform surrogate models in predicting subjective psilocybin experience even when controlling for spatial contiguity (p=1.00e-6, FDR-corrected for 79 steps in *n*; supplementary Figure 2).

Interestingly, the majority of explained variance in *b* of the idiosyncrasy-informed models was always due to either *y* or *z*, but not *x*, suggesting that *x* captured a heterogeneously distributed FC pattern that was not relevant to subjective psilocybin experience such as potentially scanning session differences (Figure 4C). Thus, we explored whether the inclusion of *x* as a noisy predictor might explain why we only found three slightly significant idiosyncrasy-informed linear models, namely at *n*={2250, 2500, 2750} (p=0.0499; p=0.0450; p=0.0480, respectively; Figure 4C). Indeed, repeating the analysis with idiosyncrasy-informed single-predictor models (i.e. *b*∼*x*; *b*∼*y*; *b*∼*z*) revealed that *b* is best explained by *y* at *n*=1000 (p=0.0082; R²=0.31) (Figure 4D-E). Thus, we will henceforth focus on the FC pattern described by PC2 corresponding to *y* at *n*=1000, which accounts for 9.91% of the variance in the 1000 most idiosyncratic edges. Notably, this PC2 is strongly inversely related to the FC pattern that corresponds to PC3 and *z* at *n*=6500, where *z* most significantly predicts *b* (p=0.0169; R²=0.27) (supplementary Figure 3).

The resulting subject scores *b* of the first principal component (PC1) were used as the response variable. Subsequently, additional PCAs were run on a set of *n* FC edges from all subjects. Edges were either selected randomly or based on high ICC, and the resulting subject scores (*x*, *y*, *z*) of the first three principal components (PC1-3) were used as predictor variables. (B) Coefficients of PC1 of the behavioural data PCA (11-dimensions of altered states of consciousness (Dittrich, 1998; Studerus et al., 2010): UN = Experience of Unity; SP = Spiritual Experience; BS = Blissful State; IS = Insightfulness; DB = Disembodiment; IC = Impaired Control and Cognition; AX = Anxiety; CI = Complex Imagery; EI = Elementary Imagery; SY = Audio-Visual Synesthesia; CM = Changed Meaning of Percepts). (C) R² value of linear models *b*∼*x*+*y*+*z* based on the *n* top ICC-ranked edges (red-line) and average R² value with corresponding standard error of 100 linear models based on *n* random selections of edges (grey line). Significant (p<0.05; uncorrected) ICC-ranked models are marked with red triangles. The *** indicate that the ICC-ranked model at *n*=2500 explains significantly (p=1.00e-6, FDR-corrected across the 79 steps in *n*) more variance in *b* than the models based on random edge selections. (D) R² of linear models *b*∼*x* (blue), *b*∼*y* (teal), *b*∼*z* (yellow) based on the *n* top ICC-ranked edges. Filled and empty triangles highlight significant (p<0.05; uncorrected) positive and negative correlations, respectively. (E) Correlation of *b* with *y* of the most significant idiosyncrasy-informed model *b*∼*y* at *n*=1000 edges.

### Characterising FC patterns of subjective psilocybin experience

The PC2 of the 1000 most idiosyncratic edges of the psilocybin group is characterised by strong, positive loadings on edges of the PCC and mPFC and strong negative loadings on edges of right anterior cingulate cortex (ACC) (Figure 5A).

**Figure 5.**
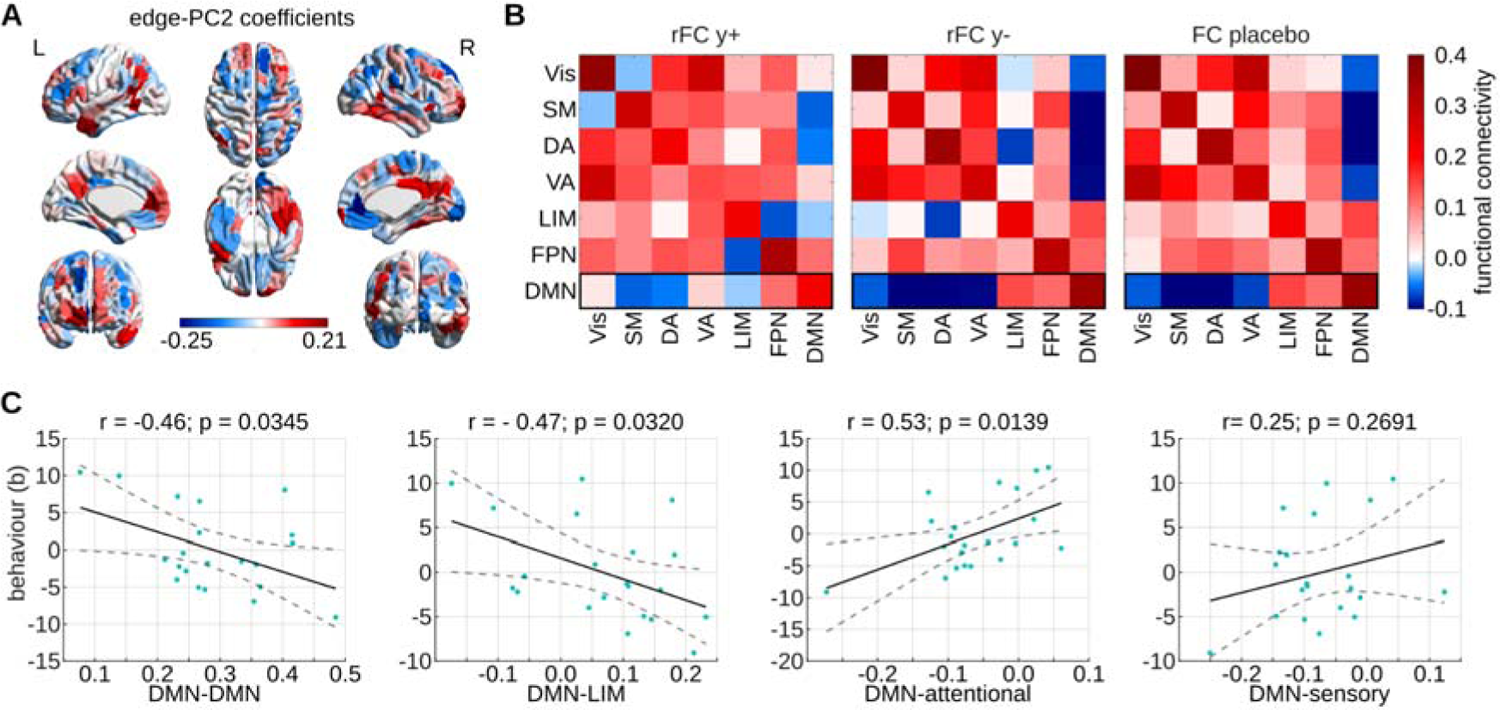
Characterisation of idiosyncratic FC pattern that predicts subjective psilocybin experience. (A) Coefficients of the PC2 corresponding to the best ICC-based model *b*∼*y* at *n*=1000. Coefficients are summed across all edges (within the subset of *n*) of each brain region to obtain brain-regional values. (B) Mean RSN functional connectivity of psilocybin subjects with positive *y* score (rFC y+) and negative *y* score (rFC y-), as well as of placebo subjects. Means are computed over the 1000 FC edges with highest ICCpsi. For all psilocybin subjects these edges are not directly derived from the FC but instead reconstructed from the edge PC2 and the corresponding subject scores *y*. Finally, the means displayed by rFC y+ and rFC y- are weighted according to the absolute *y* value of the subjects. (C) Correlations of *b* (i.e. subjective psilocybin experience) with DMN functional connectivity in psilocybin subjects. DMN functional connectivity values are averages over the 1000 edges with highest ICC in psilocybin, which were directly derived from the FC of each subject. DMN-attentional connectivity was computed as the average over DMN-VA and DMN-DA edges, and DMN-sensory connectivity corresponds to average DMN-Vis and DMN-SM connectivity. Vis = visual network; SM = somatomotor network; DA = dorsal attention network; VA = ventral attention network; LIM = limbic system; FPN = frontoparietal network; DMN = default-mode network.

To better visualise the FC pattern captured by PC2, we reconstructed the functional connectivity values of the 1000 most idiosyncratic edges for each psilocybin subject from PC2 and *y*, and averaged the reconstructed FC edges across RSN connections. This resulted in one 7×7 RSN FC matrix per subject, which were subsequently averaged across subjects with positive *y* (rFC y+), and across subjects with negative *y* (rFC y-). Averages were weighted according to the absolute value of *y* (Figure 5B). This analysis intended to crystallise the FC patterns that are representative for the “archetype” psilocybin subjects with particularly intense (rFC y+) or mild (rFC y-) psilocybin experiences (*b*), which, according to our best linear model *b*∼*y*, are associated with positive and negative values for *y,* respectively (see Figure 4E). Finally, we also computed the average RSN FC of placebo subjects based on the same (non-reconstructed) 1000 edges with maximal ICCpsi.

Intriguingly, rFC y-was strikingly similar to the placebo pendant, whereas rFC y+ was distinct (Figure 5B). Notably, this trend was particularly pronounced with respect to DMN connectivity: while the DMN connectivity of rFC y-was nearly identical to placebo, rFC y+ appeared to have reduced within-DMN and DMN-LIM connectivity and increased connectivity between DMN and attentional (DA, VA), and sensory (Vis, SM) networks. The similarity of rFC y-but not FC y+ with the placebo counterpart is particularly interesting given the interpretation of *b* in terms of the intensity of subjective psilocybin experience (Figure 4B). In other words, psilocybin subjects with negative *y* are predicted to have less intensely felt psilocybin experiences and thus, it seems unsurprising that their isolated FC pattern resembles that of placebo subjects.

Corroborating the above suppositions, we find that, with the exception of average DMN-sensory connectivity (r=0.25, p=0.2691, fdr=1.0000), average DMN-DMN (r=-0.46, p=0.0345, fdr=0.0498), DMN-LIM (r=-0.47, p=0.0320, fdr=0.0498), and DMN-attentional (r=-0.53, p=0.0139, fdr=0.0498) connectivity are significantly correlated to *b* (Figure 5C). (Note: DMN-sensory connectivity refers to the average across DMN-Vis and DMN-SM edges. Similarly, DMN-attentional connectivity is average DMN-VA and DMN-DA connectivity.) All significant correlations survive FDR correction. Importantly, if edges are not pre-selected based on idiosyncrasy and the above correlations are instead computed across all FC edges, only DMN-LIM connectivity is significantly correlated to *b* after FDR correction (supplementary Figure 4). This further supports the argument that idiosyncrasy represents a useful edge-selection criterion for improving predictions of subjective psilocybin experience, and possibly behaviour in general.

## Discussion

Human neuroimaging studies have traditionally focused on characterising populational-level blueprints of brain activity and structure by averaging over data from many subjects (Finn et al., 2015). While this has led to important insights into the general principles of neuroscience, there is no doubt that substantial variability exists not only between but also within populations. Notably, a stronger research focus on idiosyncratic rather than generalisable features may be key to the advancement of personalised medical approaches. Arguably, it is especially important in the context of psychedelic neuroimaging studies, as psychedelic drugs hold great therapeutic promise while at the same time they are known to induce highly variable responses in individuals (Moujaes et al., 2022).

Our study contributes to this end by demonstrating that idiosyncrasy can serve as a useful heuristic for identifying clinically relevant brain connectivity patterns that map onto individual drug responses. Additionally, we presented a simple, brain-fingerprinting-inspired framework for isolating idiosyncratic FC patterns. Applying this framework to high-resolution fMRI data of healthy volunteers under the acute influence of psilocybin or a placebo, we revealed that psilocybin appears to alter not only *how* identifiable subjects are based on their FC, but also *what* makes them identifiable - that is to say, which features in the FC are most idiosyncratic. Finally, we were able to extract idiosyncratic functional connections between cortical RSNs that predicted the unique drug experience of individual psilocybin subjects.

### Psilocybin-induced changes to the functional connectome fingerprint

Whereas psilocybin did not seem to affect the temporal stability (*Iself*) of FC edges, our results implied significantly greater FC heterogeneity (lower *Iothers*) in the psilocybin than in the placebo group. This finding sheds an interesting new light onto the effects of psilocybin on BOLD signal dynamics. Specifically, despite previous work reporting that acute psilocybin leads to increased BOLD signal variability (Tagliazucchi et al., 2014) and the formation of short-lived communities of interacting brain regions (Petri et al., 2014), our finding suggests that an individual’s FC nevertheless exhibits unique features that remain stable over the course of a few minutes, in line with previous work (Van De Ville et al., 2021). This is relevant because, although the subjective effects of psilocybin have been described as “labile” (Griffiths et al., 2006), these effects are arguably unlikely to rapidly shift on a minute-by-minute basis. Our findings suggest that the stable and unique temporal features carried forward in the FC can be informative about the subjective effects of psilocybin. Notably, this highlights the utility of assessing the reliability of FC features in terms of temporal stability.

Additionally, the finding of lower Iothers and unchanged Iself in the psilocybin versus placebo group also matches the intuition that the subjective experiences of individuals, who are lying in an fMRI scanner, are likely to be more heterogeneous when under the influence of peak psilocybin drug effects. Indeed, as aforementioned, psychedelic experiences are commonly characterised by fluctuations across various cognitive and emotional aspects (Girn et al., 2020; Griffiths et al., 2006). Remarkably, similar work mapping functional connectome fingerprints of a group of individuals at baseline and after administration with ayahuasca, a psychedelic compound that targets the same 5-HT2AR neuroreceptor as psilocybin, observed a decrease in FC heterogeneity (higher *Iothers*) in the drug condition (Mallaroni et al., 2022). While, on the face of it, this may seem to contradict our findings, it is important to note that the participants of the mentioned ayahuasca study were in fact members of the same religious group, who convene on a regular basis to practise a ritualistic ceremony that involves the intake of ayahuasca. Thus, ayahuasca may have induced comparatively homogeneous subjective experiences in the religious practitioners by triggering a shared memory of common rituals, which in turn may have been reflected by greater FC similarity. In contrast, the subjects of our study did not engage in common ritualistic practices that involved psilocybin intake and, hence, they likely had more heterogeneous associations with the psilocybin-induced psychedelic state. As such, both studies provide evidence to believe that FCs capture neural activity that is relevant to the subjective experience of acute psychedelic drug effects.

When mapping the cerebral distribution of idiosyncrasy we observed, in line with previous work (Finn et al., 2015; Mallaroni et al., 2022), that FPN regions tended to have exceptionally idiosyncratic functional connectivity profiles under non-psychedelic (placebo) conditions. Additionally, we found that in psilocybin subjects the distribution of idiosyncrasy was systematically shifted, and in particular shifted away from FPN and towards DMN and Vis regions.

Intriguingly, DMN and Vis regions are known to exhibit exceptionally high densities of the 5-HT2A receptor, which is targeted by psilocybin and believed to be the primary mediator of the drug’s hallucinogenic effects (Beliveau et al., 2017; Nichols, 2016). Moreover, both the DMN and Vis network have previously been directly associated with the subjective effects of psychedelics (Ballentine et al., 2022; R. L. Carhart-Harris et al., 2012). For instance, the DMN is assumed to play a key role in self-consciousness, and its perturbation through drugs or invasive electrical stimulation has been linked to distorted or weakened perceptions of self, commonly referred to as “ego dissolution” (Blanke et al., 2002; R. L. Carhart-Harris & Friston, 2010). Furthermore, a recent study from Ballentine and colleagues, analysing 6850 free-form testimonials of hallucinogenic experiences of various compounds, discovered a significant association between reports of drug-induced visualisations and the distinct neuroreceptor profile of cortical areas in the Vis system (Ballentine et al., 2022).

Hence, the first part of our analysis acted as an important proof of concept, indicating that a brain-fingerprinting approach might be capable of revealing the particular FC features that are relevant to the subjective experience of psilocybin.

Another noteworthy insight from the work of Ballentine and colleagues (Ballentine et al., 2022) was the finding that the subjective effects of psychedelics seem to emerge from the complex interplay of multiple neuroreceptors throughout the cortex, not limited to 5-HT2AR alone, which has hitherto been the main focus of psychedelic research. Thus, it would be highly interesting for future work to investigate the extent to which inter-individual variability in the subjective effects of psychedelics is driven by inter-individual variability in neuroreceptor expression as captured by “neuroreceptor fingerprints”.

### Functional-connectome correlates of subjective psilocybin experience

Leveraging our brain-fingerprinting-inspired framework, we isolated an idiosyncratic FC pattern that predicted subjective psilocybin experience and was characterised by altered DMN connectivity. Specifically, decreased DMN-DMN and DMN-LIM, and increased DMN-attentional connectivity was associated with an intense psychedelic drug experience, and vice versa. These results contribute to the growing body of evidence that points to a central role of DMN connectivity in psychedelic action (Gattuso et al., 2022).

Most notably, DMN disintegration has been consistently associated with the acute psychedelic state across various neuroimaging modalities and psychedelic compounds (R. L. Carhart-Harris et al., 2012, 2016; Müller et al., 2018; Muthukumaraswamy et al., 2013; Preller et al., 2018), and a recent study demonstrated that the degree of DMN disintegration in subjects treated with psilocybin was determined by their psilocybin plasma concentration (Madsen et al., 2021). Moreover, within-DMN connectivity was repeatedly shown to be inversely related to the intensity of subjective psychedelic effects (R. L. Carhart-Harris et al., 2016; Madsen et al., 2021; Smigielski et al., 2019). In particular, lower within-DMN connectivity in response to psychedelic-drug administration tends to coincide with stronger subjective ratings of ego dissolution (R. L. Carhart-Harris et al., 2016; Smigielski et al., 2019). Interestingly, while our measure of subjective psilocybin experience constituted a 1-dimensional summary of 11 aspects of psilocybin experience, it is worth mentioning that the three most dominant aspects (insightfulness, experience of unity, blissful state) were in fact subscales of the 5D-ASC dimension of “oceanic boundlessness”, commonly used to measure ego dissolution (Studerus et al., 2010).

It is largely agreed that the DMN relies on mnemonic and emotional input from the limbic system for a number of tasks, including self-referential thinking (R. L. Carhart-Harris & Friston, 2010; Lebedev et al., 2015; Mason et al., 2020). As such, it has been hypothesised that the acute breakdown of DMN-LIM coupling in response to psychedelics may prevent the DMN from accessing autobiographical memory that is essential for generating self-consciousness (Millière et al., 2018). This might explain our finding that DMN-LIM decoupling predicted intense psychedelic experiences. Remarkably, DMN connectivity with hippocampal regions of the limbic system has been reported to decrease acutely post administration with psychedelics, which was associated with subjective reports of ego dissolution (R. L. Carhart-Harris et al., 2014; Lebedev et al., 2015; Mason et al., 2020). Interestingly, a recent analysis of the dataset studied here found that in psilocybin subjects lower hippocampal glutamate levels were related to positively experienced ego dissolution, whereas elevated glutamate levels in the mPFC, a key DMN area, were linked to negatively experienced ego dissolution (Mason et al., 2020). Another limbic region that was shown to be acutely affected by psilocybin is the ACC (R. L. Carhart-Harris et al., 2012; Tagliazucchi et al., 2014), which in our study was found to have an exceptionally idiosyncratic connectivity profile. Specifically, the temporal BOLD signal variance in the ACC has been reported to increase in response to psilocybin administration (Tagliazucchi et al., 2014) and psilocybin-induced changes in the ACC BOLD signal amplitude were shown to be negatively correlated with the intensity of subjective effects (R. L. Carhart-Harris et al., 2012).

Furthermore, the Relaxed Beliefs Under pSychedelics (REBUS) theory (R. L. Carhart-Harris & Friston, 2019) offers a potential explanation for our finding that subjective psilocybin experience was correlated with DMN-attentional connectivity. This theory adopts the popular view of the brain as a hierarchical prediction machine, where top-down prior beliefs from higher-level brain regions suppress expected activity in lower-level brain regions, which in turn report-back prediction errors via bottom-up information flow (Clark, 2013). Crucially, DMN regions are believed to be situated at the top end of this hierarchy, where they orchestrate the computation of high-level priors that purportedly constrain our perception of reality (R. L. Carhart-Harris & Friston, 2019; Margulies et al., 2016). Hence, by disrupting DMN activity, psychedelics are proposed to weaken the suppressive influence of top-down priors on lower-level systems, leading to increased bottom-up signalling and, consequently, enabling unconstrained cognition such as manifest in perceptual distortions and subjectively experienced increases in insightfulness, as frequently reported by individuals in the psychedelic state (R. L. Carhart-Harris & Friston, 2019; Studerus et al., 2010). (As an interesting side note, a recent study indicated that there is a mismatch between *subjective* and *objective* ratings of insightfulness during acute stages of psilocybin (Mason et al., 2021).)

In this context, the attention system is commonly assumed to sit at an intermediate level in the hierarchy, passing up information about salient sensory stimuli to higher-level networks, including the DMN (R. L. Carhart-Harris & Friston, 2010; Margulies et al., 2016). Thus, our findings might reflect increased bottom-up signalling from attention networks to the DMN during peak psilocybin drug effects, consistent with the REBUS theory. Notably, our finding is also consistent with several lines of previous work, reporting an increase in DMN-attentional connectivity in response to psychedelics (R. L. Carhart-Harris et al., 2013b, 2016; Kometer et al., 2015; Müller et al., 2018).

We acknowledge that our results may be equally well accounted for by alternative, possibly complementary, theories that also frame psychedelic action in terms of disrupted higher-level association-network activity and increased bottom-up signalling, namely the cortico-claustro-cortical (Doss et al., 2021) and the cortico-striato-thalamocortical (Vollenweider & Geyer, 2001) theory, respectively. Yet, the fact that the REBUS model links psychedelic experience explicitly to modulations of DMN activity and connectivity makes this model especially illuminating in regards to our findings.

### Paving the way to personalised psychedelic therapy

Much of the recent scientific interest in psychedelics is attributable to their putatively remarkable therapeutic potential (R. L. Carhart-Harris & Goodwin, 2017). For instance, early clinical trials testing the antidepressant effect of psilocybin-assisted psychotherapy have yielded promising results (R. Carhart-Harris et al., 2021; R. L. Carhart-Harris et al., 2018; Davis et al., 2021; Griffiths et al., 2016; Ross et al., 2016). Nevertheless, there is strong evidence that depression is in fact a highly heterogeneous disorder, urging the call for a more personalised depression-treatment plan (Buch & Liston, 2021; Fried & Nesse, 2015). One attractive route involves the use of computational brain simulations that can be informed by patient-individual neuroimaging data (Deco et al., 2018; Deco & Kringelbach, 2014; Moujaes et al., 2022; Vohryzek et al., 2023). Specifically, future personalised depression treatments may be chosen based on a two-step approach: i) simulating the patients individual neural response to a (possibly psychedelic) treatment, and ii) predicting the treatment outcome based on the simulated neuroimaging data. Tantalisingly, this approach would enable individualised predictions of the counterfactual effects of various therapeutic interventions. In the context of psilocybin-assisted depression therapy, our study may provide key insights that can be used to inform the treatment-outcome predictions in step (ii) of the proposed approach. Indeed, several lines of clinical evidence imply that the acute psychedelic experience and the long-term antidepressant effects of psilocybin-assisted psychotherapy for depression may be coupled (Griffiths et al., 2016; Roseman et al., 2018; Ross et al., 2016). Moreover, similar results have been reported in two independent samples of patients with substance-use disorder (Bogenschutz et al., 2015; Johnson et al., 2017).

### Limitations

Our study is subject to several limitations that future work may want to address. For instance, unlike classical brain-fingerprinting approaches, where reliability of FC features is commonly measured based on test-retest data from two or more separate scanning sessions, we assessed the reliability of FC features in terms of their temporal stability across one continuous scan. While we argue that the high interpretability of our results and the fact that they are largely consistent with previous work clearly suggest that our approach provided meaningful results, it would be interesting to replicate our study with test-retest data that was acquired during separate scanning sessions. In fact, our finding that the PC1 components, resulting from the FC-edge PCAs, consistently failed to explain the subjective effects of psilocybin implies that scanning-session differences may have been responsible for the largest proportion of inter-individual FC variance.

Additionally, it may be worth noting that the choice of a between-subjects design in our study, as opposed to the perhaps more commonly employed within-subjects design in the literature, was based on two main considerations. Firstly, psilocybin is believed to induce neuroplastic changes in the brain that can have sustained effects on mood and cognition, lasting weeks or even months (R. L. Carhart-Harris & Goodwin, 2017). Such prolonged effects can confound the acute measurements in a within-subjects, but not in a between-subjects design. Secondly, the between-subjects design has been shown to reduce de-blinding risks compared to the within-subjects design, where all participants receive both the active and control condition (Aday et al., 2022; Muthukumaraswamy et al., 2021).

Finally, we detected slightly more head motion in the psilocybin group, which may have biassed the identifiability results. However, we addressed this limitation by controlling for motion when testing for group differences. Also, we note that the majority of our analysis was based on ICC, which is calculated based on the whole group of subjects. Thus, while ICC group-comparisons may have been affected by motion differences, the ICC ranks of single edges within one group are likely insensitive to motion outliers.

## Conclusion

In summary, our work points to a key role of DMN functional connectivity to regions of the default-mode, limbic, and attentional networks, in mediating the acute subjective effects of psilocybin. Additionally, our results imply that idiosyncrasy provides a useful heuristic for identifying clinically relevant FC patterns that predict individual drug responses, and we present a simple framework for isolating these patterns. Notably, this framework may prove fruitful in future pharmacological neuroimaging studies. Overall, our study chimes in with the trending call (Moujaes et al., 2022) for a stronger research focus on idiosyncratic rather than merely generalisable brain-activity patterns, paving the way for more effective, personalised medical approaches.

## Methods

### Participants

Data from 60 healthy volunteers with previous, but not recent (i.e. not within the 3 months prior to the study), experience with a psychedelic drug was collected between July 2017 and June 2018 at Maastricht University. Following a randomised, placebo-controlled, double-blind parallel group design, participants were assigned to one of two conditions (0.17 mg/kg psilocybin, or placebo) such that groups were matched for age, sex, and educational level. In the psilocybin condition, subjects were administered psilocybin dissolved in bitter lemon. In the placebo condition, subjects were administered plain bitter lemon instead. The study obtained ethical approval from the Maastricht University’s Medical Ethics committee and was in accordance with the Medical Research Involving Human Subjects Act (WMO) as well as with the code of ethics on human experimentation from the declaration of Helsinki (1964) and its amendments made in October 2013 in Fortaleza, Brazil. All participants were made fully aware of all procedures, possible adverse and expected beneficial effects, as well as their responsibilities and rights, including the right for voluntary termination without consequences. The data was collected as part of a larger clinical trial (Netherlands Trial Register: NTR6505). More detailed descriptions can be found in previous publications (Mason et al., 2020, 2021).

Out of the 60 participants, only 49 provided resting-state fMRI data. Furthermore, one subject was excluded because of missing values in the BOLD time series of five brain regions. Another subject had missing values in the BOLD time series of only one brain region in the visual subnetwork. This subject was not excluded. Instead, the corrupted time series was replaced by the average time series of the remaining 28 parcels in the visual subnetwork. Two further participants were excluded due to high head motion (more than 10% of fMRI volumes contained motion artefacts). The final sample comprised 21 psilocybin and 25 placebo subjects.

### Neuroimaging

Participants underwent structural MRI 50 min post psilocybin/placebo administration, in addition to 6-min resting-state fMRI 102 min post psilocybin/placebo administration during the peak subjective drug effects. All images were acquired in a MAGNETOM 7 Tesla MRI scanner. The following acquisition parameters were used for the fMRI scans. TR = 1400 ms; TE = 21 ms; field of view = 198 mm; flip angle = 60°; oblique acquisition orientation; interleaved slice acquisition; 72 slices; slice thickness = 1.5 mm; voxel size = 1.5 mm, isotropic. Furthermore, participants were presented with a black fixation cross on a white background during the scanning session and were asked to focus on the cross, clear their minds and lie as still as possible.

### Data preprocessing

The neuroimaging data was preprocessed following the pipeline from (Amico et al., 2017) and using the open-source MATLAB toolbox Apéro (https://github.com/juancarlosfarah/apero), which combines functionality from various software packages, including the FMRIB Software Library FSL (Jenkinson et al., 2012). We preprocessed each subject’s data only once, which involved preprocessing the entire, unsplit time series.

The T1-weighted (T1w) structural MRI images were denoised, skull stripped (HD-BET (Isensee et al., 2019)) and segmented (FSL *fast*) to obtain individual grey-matter, white-matter, and cerebrospinal fluid tissue masks for each subject. Next, the T1w images were registered to Montreal Neurological Institute (MNI-152) standard space by sequentially applying rigid-body (6 degrees of freedom (dof); FSL *flirt*), affine (12 dof; FSL *flirt*), and nonlinear (12 dof; FSL *fnirt*) transformations. The inverse transformation matrices from this latter step were used to convert the (Schaefer et al., 2018) parcellation with 200 cortical regions of interest (ROIs) from MNI-152 standard space to subject space. The parcellation was chosen based on a recent study showing that the generated connectomes were relatively robust to changes in the preprocessing pipeline (Luppi & Stamatakis, 2021). Individual grey-matter parcellations were subsequently obtained by applying the subject’s grey-matter tissue mask to the subject-space parcellation.

The fMRI BOLD time series for each ROI were extracted as follows. First, fMRI volumes were corrected for slice-timing (FSL *slicetimer*) and the skull was removed (FSL *bet*) prior to correcting for motion (FSL *mcflirt*). The signal was normalised to mode 1000 and subsequently demeaned and linearly detrended. To enable the isolation of signals from different brain tissues and ROIs, the tissue masks and grey-matter parcellations, which were previously derived from the T1w images, were registered to each subject’s fMRI mean volume via 6-dof rigid-body transformation (FSL *flirt*), followed by inverse boundary-based T1w-to-fMRI image registration (FSL *flirt-bbr*). Then, a total of 18 regressors, comprising 3 translations (x, y, z), 3 rotations (pitch, yaw, roll), the average cerebrospinal-fluid, white-matter and whole-brain (i.e. global signal regression) signals, and all corresponding derivatives, were removed from the overall BOLD signal. The resulting signal was bandpass filtered, retaining frequencies between 0.01 and 0.25 Hz, by applying a first-order Butterworth filter. Subsequently, the first principal component (PC1) of the cerebrospinal-fluid, white-matter, and whole-brain signals were, sequentially, regressed out in an additional cleaning step (note that this step is equivalent to the CompCor method described by Behzadi and colleagues (Behzadi et al., 2007)). Specifically, these principal components were included as nuisance parameters within a general linear model for the BOLD time series data. The time series were extracted by regressing out the nuisance parameters from the BOLD time series and were averaged across adjacent voxels to obtain one time series per ROI.

### Assessing head motion

The number of fMRI volumes with motion artefacts was used to gauge the amount of head motion of each participant during the scan. Following (Amico et al., 2017), motion artefacts were detected based on three parameters: 1) Frame Displacement, which estimates the displacement of the head between consecutive volumes in mm (FSL *fsl_motion_outliers*); 2) DVARS, which detects abrupt changes in signal intensity from one volume to the next and is defined as the root mean square variance over voxels of the temporal derivatives of the time series (FSL *fsl_motion_outliers*); and 3) the standard deviation of the BOLD signal across voxels at each time point. Specifically, a volume was marked to contain motion artefacts if any of the three conditions was met: 1) Frame Displacement > 0.55; 2) DVARS > 75 percentile + 1.5 of the interquartile range; 3) standard deviation > 75 percentile + 1.5 of the interquartile range. There was a slightly significant difference in motion between the groups, indicating that psilocybin subjects tended to move more than placebo subjects (mean difference=4.22; t-stat=2.06; p=0.0456). Thus, we accounted for motion as confounder wherever applicable, as described in the subsection “Statistical analysis”.

### Ratings of subjective drug experience

Participants were asked to retrospectively report their subjective drug experience by completing the “five dimensions of altered states of consciousness” (5D-ASC) questionnaire (Dittrich, 1998) 360 min post drug administration. This well-established questionnaire was originally designed to measure five aetiology-independent dimensions of the subjective experience of altered states of consciousness (ASCs). However, more recent evidence suggests that the proposed dimensions can be decomposed into eleven subdimensions with greater interpretability and improved ability to discriminate between distinct psychoactive-drug conditions (Studerus et al., 2010). Thus, our analysis focuses on these eleven subdimensions, which are: UN = Experience of Unity; SP = Spiritual Experience; BS = Blissful State; IS = Insightfulness; DB = Disembodiment; IC = Impaired Control and Cognition; AX = Anxiety; CI = Complex Imagery; EI = Elementary Imagery; SY = Audio-Visual Synesthesia; CM = Changed Meaning of Percepts. For more details, we refer the reader to (Studerus et al., 2010).

### Functional connectomes

We constructed FCs with N = 200 nodes, where each node *i* represented one of 200 cortical brain regions of the Schaefer parcellation (Schaefer et al., 2018) and was connected to every other node *j* according to the Pearson correlation of the fMRI BOLD time series of *i* and *j*. For each subject, we derived one FC from the first and another from the second half of the fMRI time series. The resulting two FCs per subject were referred to as first-half and second-half FC, respectively, and we used them to calculate the brain-fingerprinting metrics (*Iself*, *Iothers*, *Idiff*) and edgewise ICC values. Additionally, we computed another FC per subject from the unsplitted fMRI time series. Edges from this latter FC served as potential predictors of subjective drug experience.

### Brain fingerprinting

Building on the work of Amico and Goñi (Amico & Goñi, 2018), we measured the identifiability of a subject i given a set of FC edges X using the metric *Idiff*, which is defined as the difference of that subject’s *Iself* and *Iothers*.

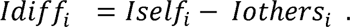

*Iself* and *Iothers* in turn can be derived from the (non-symmetric, square) identifiability matrix A, in which each element a_i,j_ encodes the Pearson correlation coefficient of the second-half FC of subject i and the first-half FC of subject j, considering the edge selection X. The diagonal elements of A correspond to the first-half versus second-half similarities of the subjects with respect to X, which we refer to as *Iself*.

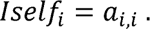

Furthermore, the off-diagonal elements of A encode the between-subject similarities with respect to X. Hence, the *Iothers* of subject *i* was calculated by averaging across the off-diagonal elements of the i-the row and column in *A*. (Note: we derived a separate identifiability matrix for each condition such that only within-group between-subject correlation values were used to compute *Iothers*.)

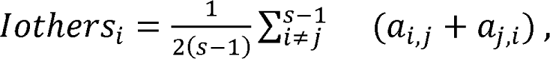

where *s* is the number of subjects in the condition of subject *i*. Taken together, edge selections with high average LJLJLJLJLJ tend to be stable throughout the six-minute fMRI scan of the same subject, but vary strongly between subjects. In other words, they are highly subject-specific just like a person’s individual fingerprint.

### Intraclass correlation

Intraclass correlation (ICC) represents a widely used statistical measure to quantify the reliability of repeated measurements taken from data that is structured in groups. A high ICC value indicates high similarity among measurements from the same group and strong between-group variance. Importantly, previous work on brain fingerprinting using ICC to assess the reliability of FC traits (= groups) across test-retest FCs (= measurements) suggests that selecting edges based on high ICC can improve subject identifiability and behavioural predictions (Amico & Goñi, 2018; Finn et al., 2015). Following this rationale, we calculated class-1 intraclass correlation of FC edges across first- and second-half FCs as follows.

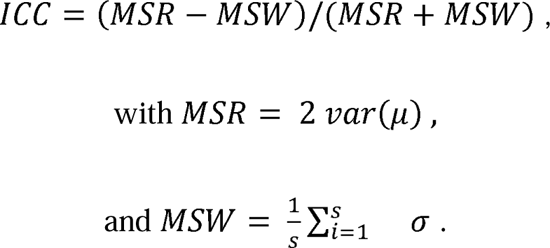

Here, var(µ) is the variance of LJ, LJ is a vector encoding the average edge value across first- and second-half FCs for each subject, s is the number of subjects, and LJ is a vector encoding the variance of edge values across first- and second-half FCs for each subject. Put simply, MSR is the variance of group means and MSW is the total within-group variance. Hence, an FC edge with high ICC tends to differ strongly between subjects, while it remains temporally stable throughout the scan of the same subject. (Note: we computed the edgewise ICCs for each condition separately as ICC is blind to potential systematic differences between groups of subjects.)

### Predicting subjective psilocybin experience

To examine whether idiosyncratic FC patterns allow for improved predictions of inter-individual differences in subjective psilocybin experience, we leveraged PCA to isolate independent, heterogeneously distributed traits and reduce the dimensionality of our prediction target (subjective psilocybin experience) and the predictors (FC edges). PCA is a widely applied statistical technique that can be used to re-represent the data in the space of principal components (PCs). Importantly, these PCs are orthogonal and hierarchically ordered such that the first and last PCs point in the direction of the largest and smallest variance in the data, respectively. Hence, each PC accounts for an independent proportion of variance in the data, which is encoded in the projections of each data point onto the PC, a.k.a. the PC scores.

We ran PCAs on two different sets of data: i) the behavioural data (21 x 11; subjects x ASC aspects), and ii) the functional connectivity values of *n* FC edges from all subjects (21 x n; subjects x FC edges), where the *n* edges were either selected based on idiosyncrasy (ICCpsi) or randomly. The resulting PCs can therefore, respectively, be interpreted in terms of independently distributed ASC and FC patterns with the corresponding PC scores indicating the extent to which these patterns were present in each subject. Subsequent to the PCAs, we constructed linear models to predict subjective psilocybin experience from FC edges, where the former was represented by the scores of PC1 from the behavioural PCA and the latter by the scores of PC1-3 from the FC-edge PCA.

The choice of our prediction-target representation was based on the fact that the PC1 of the behavioural PCA accounted for a reasonable proportion of variance in the data (45.7%) and it captured a highly interpretable ASC pattern that is arguably of great interest, namely the intensity of the drug experience (see Fig. 4b).

The line of reasoning behind the construction of our predictors shall be illustrated in the following. Firstly, our motivation for using PCA in this context was to remove potential redundancies in the selected edges and isolate the independent, maximally heterogeneous FC patterns that we are interested in. Furthermore, recall that the core hypothesis of our study is that selecting edges based on idiosyncrasy has the effect of enriching for inter-individual variance that maps onto heterogeneous behavioural traits. That is, behaviourally relevant neural processes are hypothesised to be more important contributors to the variance of idiosyncratic edges than to that of random edges. As such, we expect that in idiosyncratic edges, behavioural heterogeneity may be reflected by one or more of the few most important PCs, whereas in random edges it may be reflected by less important PCs, or not at all. For this reason, we argue that a small number of PC-score predictors *k* is sufficient for testing our hypothesis. Specifically, evidence in support of our hypothesis would be if idiosyncrasy-informed models consistently outperform random models for a constant, small *k*, and the vice versa would represent conflicting evidence. We used *k*=3, however we show that our overall results are replicated with different choices of *k* (supplementary Fig. 5). We further argue that choosing a small number for *k* is not only sufficient but also preferable to avoid the risk of overfitting as demonstrated in supplementary Fig. 5, showing that with *k*=20 all idiosyncrasy-informed and random models explain 100% of subjective psilocybin experience. Finally, although in classical PCA-based predictive modelling approaches it is common to choose *k* such that the cumulative percentage of variance in the data explained by PC1-*k* is close to 100%, this rule of thumb was not reasonable in our case i) because of the risk of overfitting, as aforementioned, and ii) because this approach would have required us to change *k* for each *n*, which was incremented at each iteration, and for each random model — even if we had done so — it would not have been possible to perfectly control the percentage of variance explained by PC1-*k*, rendering model comparisons difficult.

## Supporting information

Supplementary Information, revised

## Statistical analysis

When comparing brain fingerprinting measures between drug-placebo conditions, we included motion as a potential confounding factor in the linear model to predict the measure of interest from the categorical group variable (i.e. y ∼ group + motion), and reported the effect of the group variable. Otherwise, the default statistical test was a paired t-test. Where applicable, False Discovery Rate (FDR)-corrected p-values are given (indicated as “fdr”).

## Data availability

The connectomes and the accompanying covariates used to differentiate individuals can be made available to qualified research institutions upon reasonable request to J.G.R and a data use agreement executed with Maastricht University.

## Code availability

All code used for analysis will be made available upon acceptance on E.A.’s GitHub page (https://github.com/eamico/).

## Acknowledgements

E.A. acknowledges financial support from the SNSF Ambizione project “Fingerprinting the brain: network science to extract features of cognition, behaviour and dysfunction” (grant number PZ00P2_185716).

## Author contributions

J.G.R., and N.M. acquired the data. H.T. and J.C.F. preprocessed the data and extracted the functional connectomes. H.T. and E.A. conceptualised the study and designed the framework. H.T. performed the connectome fingerprinting analyses and wrote the first version of the manuscript. E.A. supervised the study. All authors interpreted the results and revised the manuscript.

## Competing interests

None.

## Notes

### Competing Interest Statement

The authors have declared no competing interest.

### Summary of Updates

Revised version.

