## Supplementary Information, revised for "The Unique Neural Signature of Your Trip: Functional Connectome Fingerprints of Subjective Psilocybin Experience"

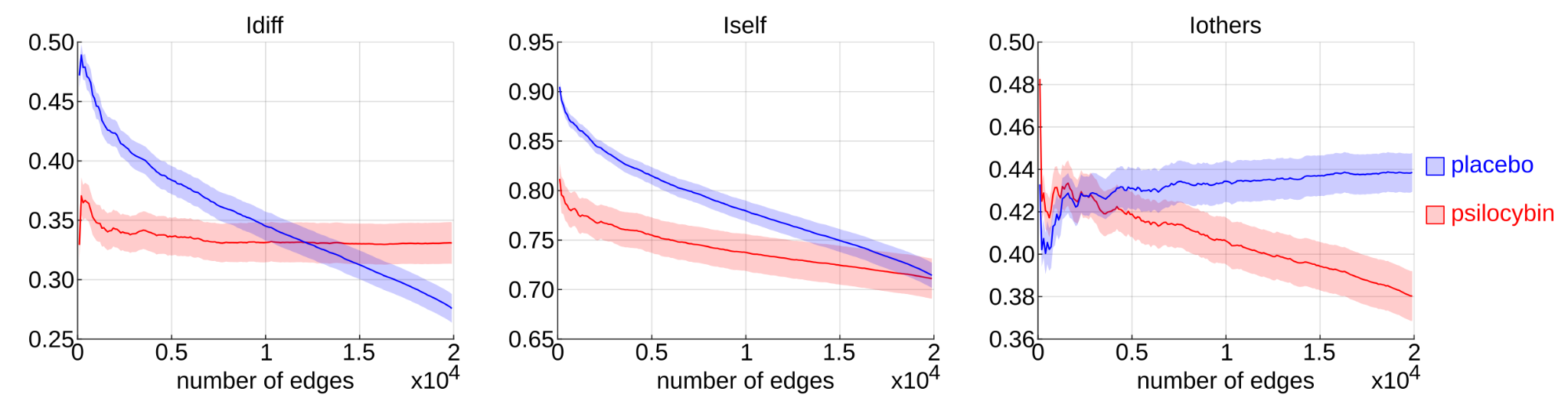

Supplementary Figure 1. Selecting edges based on high ICC in placebo optimises Idiff in the placebo but not psilocybin group. Group-average and corresponding standard error of *Idiff*, *Iself*, and *Iothers* in placebo (blue) and psilocybin (red) computed for the *n*={50, 100, …, 19900} edges with the highest ICCpla.

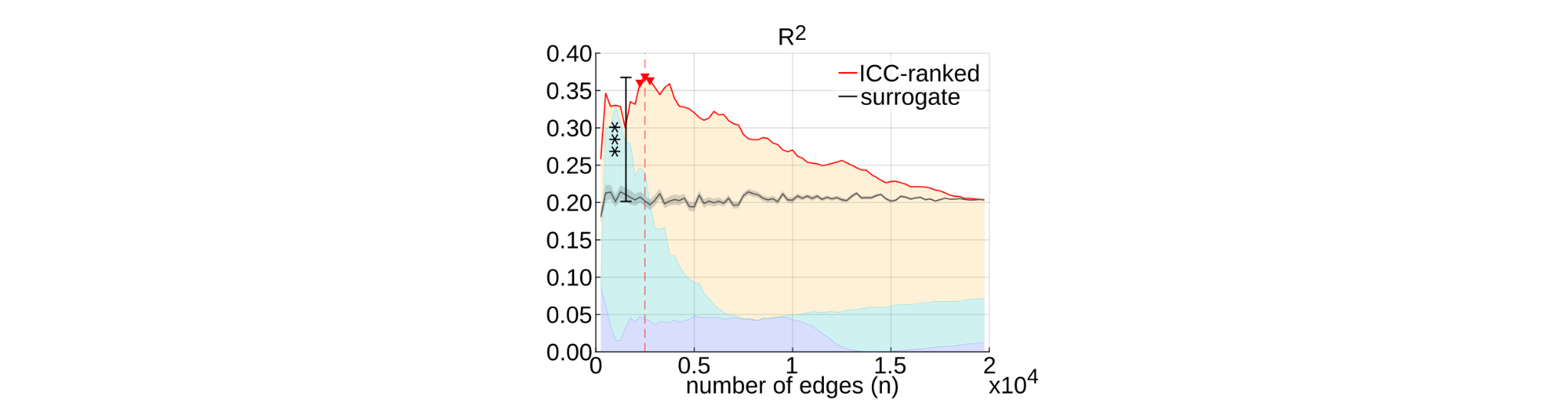

Supplementary Figure 2. R² value of linear models *b*~*x*+*y*+*z* based on the *n* top ICC-ranked edges (red-line) and average R² value with corresponding standard error of 100 surrogate linear models based on *n* edges that were selected such that spatial contiguity was preserved (grey line). Significant (p<0.05; uncorrected) ICC-ranked models are marked with red triangles. The *** indicate that the ICC-ranked model at n=2500 explains significantly (p=1.00e-6, FDR-corrected across the 79 steps in n) more variance in *b* than the spatial-contiguity-preserving models.

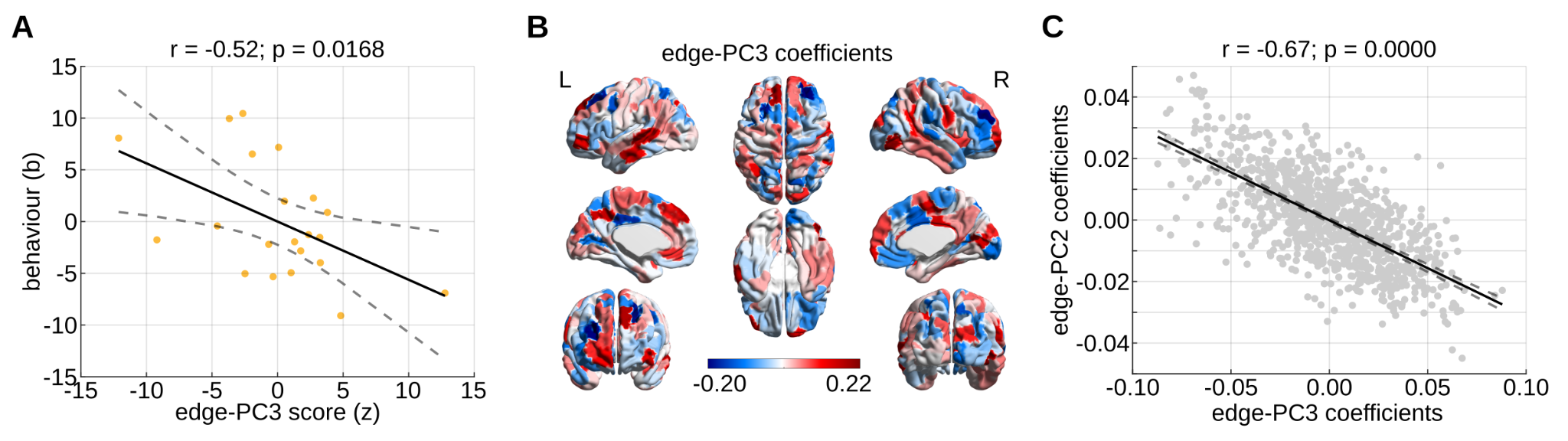

Supplementary Figure 3. Strong correspondence between the principal-component predictors of the most significant ICC-based single-predictor models *b*~*y* and *b*~*z*. (A) Correlation of *b* with *z* of the most significant ICC-based model *b*~*z* at *n*=6500 edges. Analogous to Fig 4e in the main manuscript. (B) Coefficients of the PC3 corresponding to the best ICC-based model b~z at n=6500. Coefficients are summed across all edges (within the subset of *n*) of each brain region to obtain brain-regional values. (C) Correlation between the coefficients of edge-PC2 (at *n*=1000) and the first 1000 coefficients of edge-PC3 (at *n*=6500). Note the high correspondence between the two principal components, suggesting that they are essentially capturing the same (sign-flipped) pattern of variance.

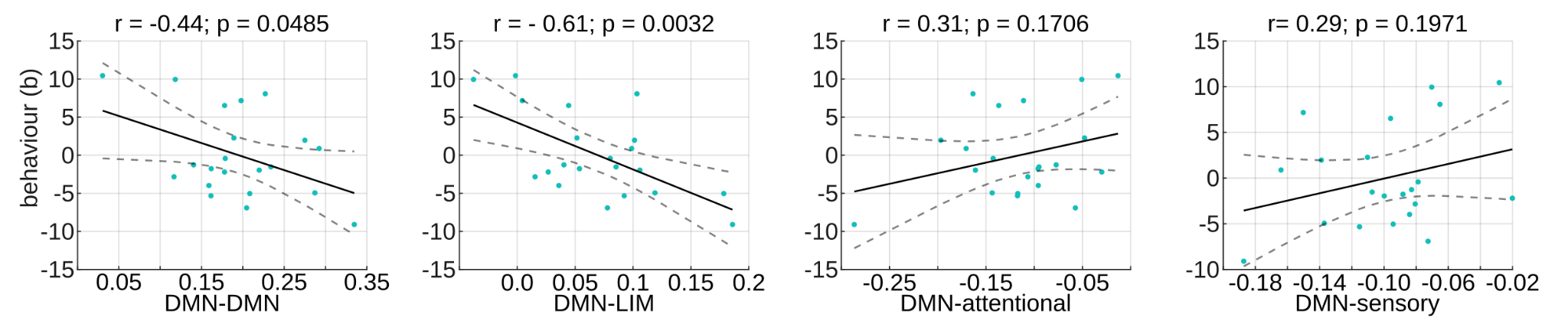

Supplementary Figure 4. Correlations of *b* (i.e. subjective psilocybin experience) with DMN functional connectivity in psilocybin subjects, where the DMN functional connectivity values are averages over all edges between all brain regions of the DMN and the LIM, attentional (VA, DA), or sensory (Vis, SM) networks. Edges were directly derived from the FC of each psilocybin subject. The given p-values are uncorrected. FDR-corrected p-values are fdr=0.1000, fdr=0.0138, fdr=1.0000, and fdr=1.0000 for behavioural correlations with DMN-DMN, DMN-LIM, DMN-attentional, and DMN-sensory connectivity, respectively. Vis = visual network; SM = somatomotor network; DA = dorsal attention network; VA = ventral attention network; LIM = limbic system; FPN = frontoparietal network; DMN = default-mode network.

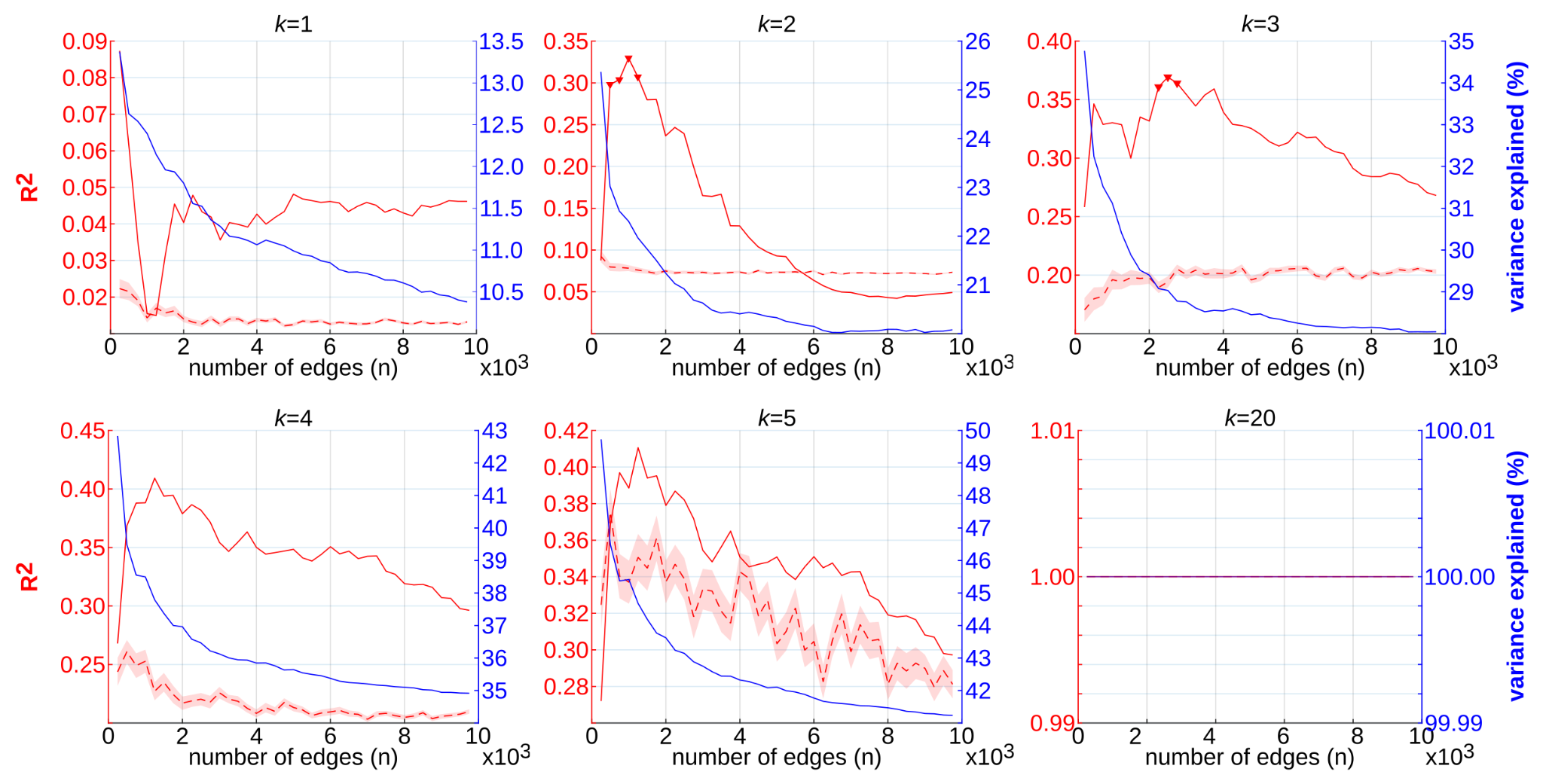

Supplementary Figure 5. R² values and percentage of explained variance of linear models to predict subjective psilocybin experience from *n* FC edges, which were either selected based on idiosyncrasy (ICCpsi) or randomly. Subjective psilocybin experience is represented in the models by the scores corresponding to the first principal component (PC) of the behavioural data (see main manuscript Fig. 4a). The model-predictor variables are the scores corresponding to the first *k* PCs of the *n* FC edges of all psilocybin subjects. Furthermore, the R² value of idiosyncrasy-informed models is plotted as a solid red line, while the dashed red line indicates the average R² value of 100 models based on different random edge selections and the corresponding standard error is given by the red transparent area. Significant (p<0.05; uncorrected) predictions of idiosyncrasy-informed models are marked with triangles. Finally, the solid blue line displays the cumulative percentage of variance in the *n* FC edges that is explained by the first *k* PCs. Note that idiosyncrasy-informed models tend to outperform random edge-selection-based models in explaining subjective psilocybin experience for small *k*. However, if *k* is large (*k*=20) all models explain 100% of subjective psilocybin experience, which is likely due to overfitting.
